# Performance of shotgun metagenomic sequencing for detection of fungi and parasites across clinical sample types: a multicenter retrospective study

**DOI:** 10.64898/2026.03.18.712591

**Authors:** Théo Ghelfenstein-Ferreira, Cécile Angebault, Vanessa Demontant, Laure Boizeau, Sandrine Houze, Christophe Rodriguez, Françoise Botterel

**Affiliations:** AP-HP, Hôpital Henri-Mondor, Unité de Parasitologie-Mycologie, Département de Prévention, Diagnostic et Traitement des Infections, F-94010 Créteil, France; Université Paris-Est Créteil, INSERM, IMRB, U955, F-94010 Créteil, France; AP-HP, Hôpital Henri-Mondor, Genobiomics platform, Département de Prévention, Diagnostic et Traitement des Infections, F-94010 Créteil, France; Université Paris Cité, UMR 261, Merit, IRD, F-75006 Paris, France; AP-HP, Hôpital Bichat, Centre National de Référence du Paludisme, F-75018 Paris, France; Université Paris-Est Créteil, AH-HP, Hôpital Henri-Mondor, Unité de Virologie, Département de Prévention, Diagnostic et Traitement des Infections, INSERM U955, Team Viruses, Hepatology, F-94010 Créteil, France

## Abstract

To evaluate the diagnostic performance of clinical shotgun metagenomic sequencing (SMg) for detecting medically relevant fungi and parasites compared with standard of care (SoC), and to define read-based thresholds for interpretation, we retrospectively analyzed 198 clinical samples from 187 patients across four university hospitals (2018–2022): blood (n=37), faeces (n=63), respiratory fluids (n=54), other biological fluids (n=24), and tissue biopsies (n=20). Total nucleic acids were sequenced (≥10 million reads per library) and processed with MetaMIC v2.2.1. Data were normalized as reads per million (RPM). Receiver operating characteristic analyses were used to derive optimal RPM thresholds by sample type. SoC identified microorganisms in 152/198 samples (76.8%). All 46 SoC-negative samples were also negative by SMg. At the genus level, SoC identified 187 taxa and SMg 175. Of these, 147 (78.6%) were detected by both methods, 40 (21.4%) by SoC only, and 28 (14.9%) by SMg only. The overall genus-level F1-score was 0.84. Quantification cycle (Cq) values (n=57) correlated inversely with RPM (p<0.001), and no false negatives occurred with Cq<28.6. Optimal thresholds were 0.06 RPM for faeces (AUC 0.89), 0.07 for respiratory fluids (AUC 0.93; sensitivity 88.9%, specificity 90.7%), 0.09 for blood (AUC 0.99), 0.19 for other fluids (AUC 0.94), and 0.57 for biopsies (AUC 0.89). A global threshold of 0.06 RPM yielded an AUC of 0.92 (sensitivity 88.9%, specificity 88.5%). A pragmatic uniform 0.1 RPM threshold maintained performance, while sample-type specific thresholds further improved accuracy, supporting standardized implementation of clinical metagenomics for fungal and parasitic diagnostics.

## INTRODUCTION

Clinical shotgun metagenomic next-generation sequencing (SMg) [1,2] presents specific challenges in diagnosing infections caused by eukaryotic pathogens, including fungi and parasites. These microorganisms have complex, robust cell walls that require specialized lysis methods, often resulting in low nucleic acid yields and reduced SMg sensitivity [3–7]. Furthermore, eukaryotes are typically present in low abundance in clinical samples, even during active infection [8–11]. This low pathogen load, combined with extraction issues, may further compromise SMg performance for fungi and parasites. The genomic similarity between eukaryotic pathogens and the human host increases the risk of misclassification during bioinformatics analysis. Host sequences may be misidentified as pathogens, or true pathogen reads may be erroneously excluded, leading to false positives and negatives [12,13]. Although guidelines and recommendations exist to standardize SMg for pathogen detection, they are primarily designed for bacterial [14,15] and viral [16–18] pathogens and cannot be applied directly to fungi and parasites. The lack of specific guidelines for eukaryotes, particularly regarding positivity thresholds and read count interpretation, highlights a critical gap in SMg. Without comprehensive studies exploring variability in read counts across diverse eukaryotic species and sample types, establishing reliable detection criteria remains highly challenging. Our study aimed to evaluate the analytical performance of SMg in detecting fungi and parasites across a wide range of clinical samples.

## MATERIALS AND METHODS

We followed the STROBE-Metagenomics guidelines [19], with full methods detailed in the supplementary materials.

### Sample selection and reference identification

A total of 198 clinical samples were collected retrospectively from adult patients at four French university hospitals (Henri-Mondor, Kremlin-Bicêtre, Paul Brousse, and Bichat Claude-Bernard) between January 1, 2018 and December 31, 2022. These were residual samples from standard-of-care (SoC) mycology and parasitology laboratory testing. For fungi, SoC included direct microscopy, culture, morphological identification, MALDI-ToF analysis using the MSI-2 platform [20], and quantitative PCR (qPCR). For parasites, the SoC included direct microscopy, concentration techniques, Novodiag® Stool Parasites assay (Mobidiag, Espoo, Finland) [21], and qPCR. Following routine analysis, the samples were stored at −20°C and manually selected to ensure broad representation of fungi and parasites, as well as diverse sample types.

### Clinical shotgun metagenomics

Each sample underwent a pre-extraction process combining mechanical (glass beads), enzymatic (proteinase K), and chemical lysis (RLT Buffer, Qiagen, Hilden, Germany). Whole nucleic acids were extracted using the QIAsymphony DSP Midi Kit (Qiagen) as previously described [22]. To monitor sequencing performance and exclude contamination, a positive control (ZymoBIOMICS Microbial Community Standard, Zymo Research, Freiburg, Germany) and an environmental control (DNA/RNA-free water) were processed in parallel with each extraction batch (Fig. S1). DNA and RNA libraries were prepared using the DNA Prep and TruSeq Total RNA Kits (Illumina, San Diego, CA), respectively. Sequencing was performed on an Illumina NovaSeq 6000 system using a 300-cycle paired-end protocol. Samples were retained for analysis if both the DNA and RNA libraries reached a minimum depth of 10 million reads each [18,23]. Reads from the DNA and RNA libraries were then pooled for further analysis.

### Bioinformatic analysis

A sample was considered acceptable for analysis if RNA and DNA sequences generated exceeded 10 million, respectively. After two consecutive attempts, if the number of sequences remained below 10 million, the sample was classified as low cellularity and still included in the analysis. If the expert consortium result was identical between the reprocessing and the initial run, the initial run was retained for analysis. The sequencing data were analyzed exclusively for medically relevant fungi and parasites using the ISO 15189-accredited, already published MetaMIC software (version 2·2·1) [17,22]. This process began with the removal of adapters and the exclusion of low-quality and low-complexity sequences. The remaining sequences (preprocessed reads) were then compared with a nucleotide-based database to identify potential pathogens. To ensure the reliability of the findings, these sequences were also compared with those from environmental controls. The analysis algorithm employed by MetaMIC is designed to accurately identify pathogens in sequencing data through a two-step final process: (i) Exclusion of human, mammalian and plant sequences; (ii) Sequence counts were made for each fungal and parasite species and compared to environmental control. Microbial read counts were normalized and expressed as reads per million (RPM). RPM were calculated by dividing the number of organism-specific reads by the total number of quality-filtered reads. For any RPM value of zero, a value of 0·0001 was added prior to log transformation to allow accurate plotting on a logarithmic scale.

### Data analysis by expert consortium

When metagenomic sequencing was the only positive result and samples were available, orthogonal testing was performed using confirmatory methods such as direct examination, qPCR, or analysis of concomitant samples. Two groups of specialists, SMg microbiologists (FB and CR, TGF and CA), analyzed the SMg results for each patient in a blinded, independent manner. In the event of disagreement between the two groups, a final consensus was reached through collective deliberation.

### Ethics statement

This non-interventional study complies with French law (April 15, 2019) and the Declaration of Helsinki. The database was registered with the French Data Protection Authority (Commission Nationale Informatique et Liberté) (no. 2240073) and with the AP-HP General Data Protection Regulation (RGPD) (no. 20250822143124). As samples were collected during routine clinical work and patient-identifiable information was anonymized during analysis, written or verbal informed consent was waived.

### Statistical analysis

The reference standard was a composite gold standard combining retrospective SoC results and orthogonal testing (Table S1). SMg detections inconsistent with SoC and not confirmed by orthogonal testing were classified as false positives; those confirmed were considered true positives. Concordance between SoC and SMg was quantified using the F₁-score [24]. The agreement categories were as follows: (i) Total agreement (F₁ = 1), the microorganisms detected by SMg matched exactly those identified by SoC; (ii) Partial agreement (0 < F₁ < 1), discrepancy in the microorganisms detected; and (iii) No agreement (F₁ = 0), SMg failed to detect any of the microorganisms identified by SoC. F₁-score was expressed as: 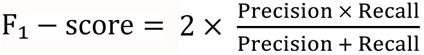 where 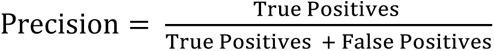 and 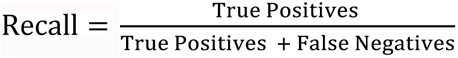.

Due to NCBI database limitations, all taxa within the Saccharomycotina subphylum (including *Saccharomyces*, *Nakaseomyces*, *Cyberlindnera*, *Pichia*, *Clavispora*, and *Candida*) were grouped together for genus-level analysis. Normality of continuous variables was assessed with the Shapiro–Wilk test. Data were expressed as the median (interquartile range, IQR). Depending on distribution, groups were compared using one-way ANOVA (with Dunnett’s multiple comparisons) or Kruskal–Wallis with Dunn’s post hoc. Two-group comparisons used the Mann–Whitney U test when non-parametric. Categorical data were compared with the Chi-2 test or Fisher’s exact test. Correlations were quantified by Spearman’s *ρ,* and a 95% confidence interval was employed. Diagnostic thresholds were derived by ROC analysis against a composite gold standard (SoC plus orthogonal testing). Two-sided p < 0·05 was considered significant. Statistical analyses were performed using R software (version 4·4·2) and GraphPad Prism (version 10·4·2).

### Data and code availability

The MetaMIC code is available at https://gitlab.com/mndebi/metamic.git. Microbial raw data are available to download (BioProject no. PRJNA1391352).

## RESULTS

### Cohort description (standard of care, SoC)

A total of 198 clinical samples from 187 patients were included (Fig. 1). The samples were categorized into five main types (blood, faeces, respiratory fluids, other biological fluids, and tissue biopsies) and subdivided into 17 subtypes (Table 1). SoC yielded positive results in 152 samples and negative results in 46 (Fig. 2(a)), identifying 44 genera and 57 species. Of the 102 fungi identified, Ascomycota predominated (n = 84), followed by Mucoromycota (n = 6), Basidiomycota (n =6), and Microsporidia (n = 6). Among the 99 parasites, protozoa were the most prevalent (n = 76), alongside 23 helminths (Table S1).

**Fig. 1.**
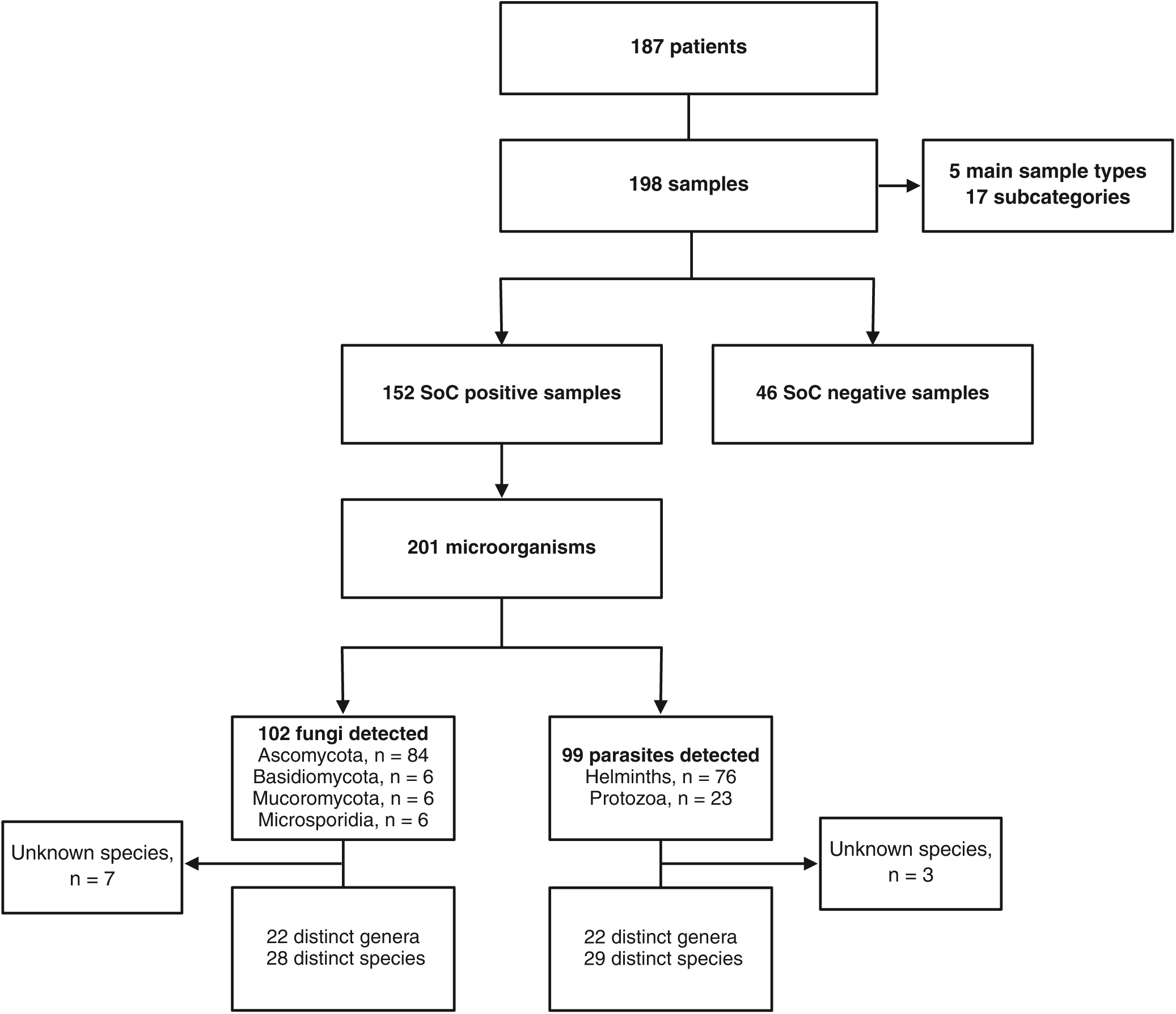
Flow chart of the study. Flow diagram illustrating the distribution of 198 clinical samples from 187 patients, classified by sample type and by standard of care (SoC) results.

**Fig. 2.**
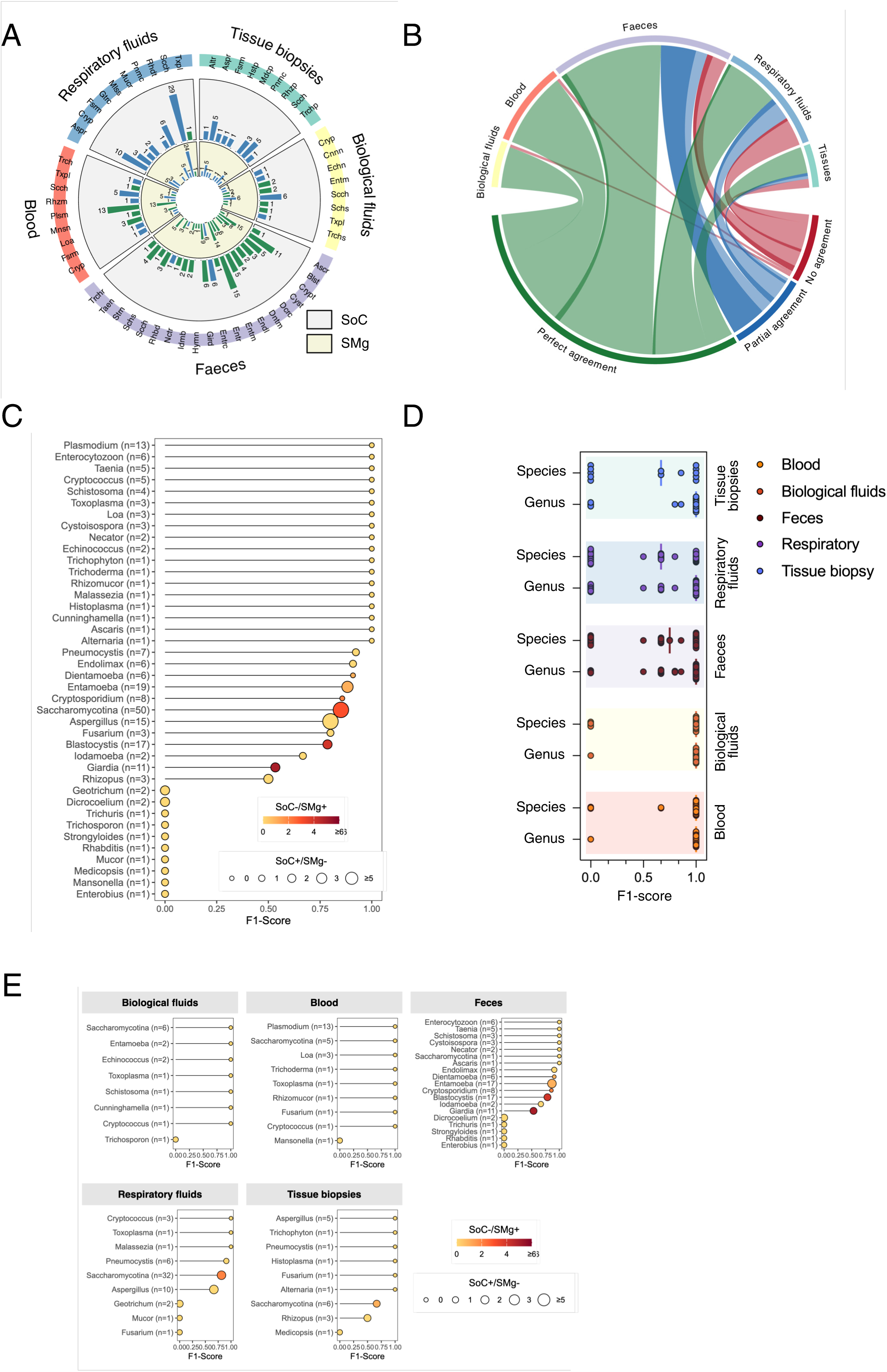
Agreement between standard of care (SoC) and shotgun metagenomics (SMg) for eukaryote detection. (a) Circular bar plot of genus-level identifications stratified by sample type (blood, respiratory fluids, tissue biopsies, biological fluids, and faeces). The outer ring represents SoC identifications (blue = fungi; green = parasites), and the inner ring represents SMg detections. (b) Chord diagram illustrating genus-level agreement across 198 samples. Links are colored by F₁-score category: perfect agreement (F₁ = 1, green), partial agreement (0 < F₁ < 1, blue), and no agreement (F₁ = 0, red). Darker links indicate additional microorganisms detected by SMg (SoC−/SMg+). (c) Global lollipop chart of genus-level F₁-scores, ordered from highest to lowest. Line length indicates the F₁-score, bubble size reflects the number of SoC+/SMg− microorganisms, and bubble color reflects the number of SoC−/SMg+ microorganisms. (d) Distribution of per-sample F₁-scores at the genus and species level according to sample type. Dots represent individual values, with medians indicated. (e) Lollipop charts of genus-level F₁-scores stratified by the five sample types, showing within-sample type variability of SMg performance. Abbreviations: Altr (*Alternaria*), Ascr (*Ascaris*), Aspr (*Aspergillus*), Blst (*Blastocystis*), Cryp (*Cryptococcus*), Crypt (*Cryptosporidium*), Cnnn (*Cunninghamella*), Cyst (*Cystoisospora*), Dcrc (*Dicrocoelium*), Dntm (*Dientamoeba*), Echn (*Echinococcus*), Endl (*Endolimax*), Entm (*Entamoeba*), Entr (*Enterobius*), Enrtc (*Enterocytozoon*), Fsrm (*Fusarium*), Gtrc (*Geotrichum*), Gird (*Giardia*), Hstp (*Histoplasma*), Hymn (*Hymenolepis*), Idmb (*Iodamoeba*), Loa (Loa), Mlss (*Malassezia*), Mnsn (*Mansonella*), Mdcp (*Medicopsis*), Mucr (*Mucor*), Nctr (*Necator*), Plsm (*Plasmodium*), Pnmc (*Pneumocystis*), Rhbd (*Rhabditis*), Rhzm (*Rhizomucor*), Rhzp (*Rhizopus*), Rhdt (*Rhodotorula*), Scch (*Saccharomycotina*), Schs (*Schistosoma*), Strn (*Strongyloides*), Taen (*Taenia*), Txpl (*Toxoplasma*), Trch (*Trichoderma*), Trchp (*Trichophyton*), Trchs (*Trichosporon*) and Trchr (*Trichuris*).

**Table 1.**
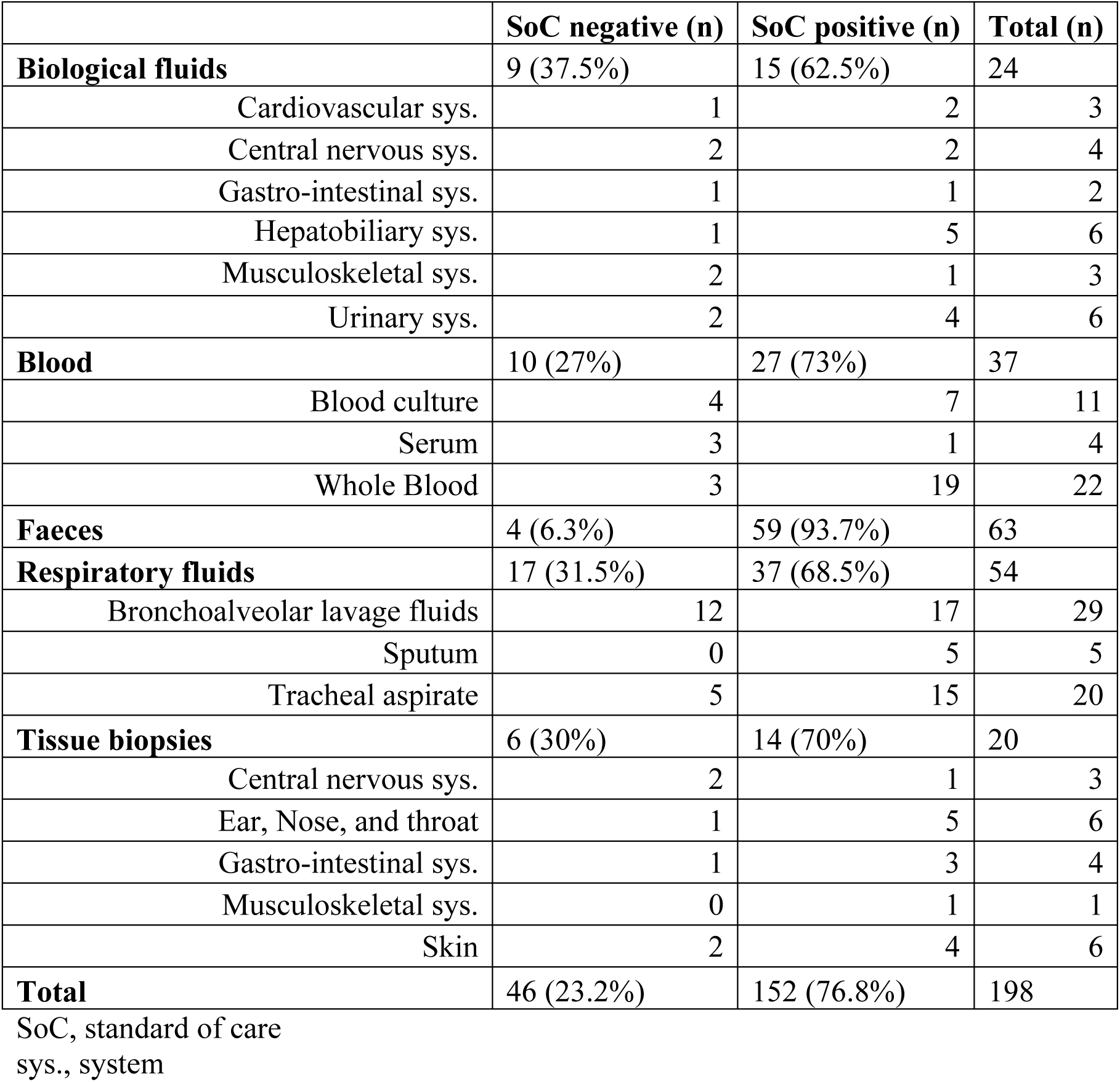
Distribution of sample types by standard of care results.

### Agreement between standard of care (SoC) and shotgun metagenomics (SMg)

The agreement between the composite gold standard (SoC plus orthogonal testing) and SMg was assessed at the genus-level using the F₁-score (Fig. 2(b)). Perfect agreement (F₁= 1) was observed for 150/198 (75·8%) samples. Of these, 104 (52·5%) were positive with SoC (SoC+), and 46 were negative (SoC−; *i.e.,* all SoC− samples). Partial agreement (0 < F₁ < 1) was observed in 26/198 samples (13·1%), mostly due SMg detecting additional microorganisms (n = 15). Complete discordance (F_1_ = 0) was observed in 22/198 samples (11.1%). SMg yielded additional detections in 21/198 samples (10·6%), predominantly in faeces (16/63, 25·4%). At the genus-level, SoC identified 187 unique taxa compared to 175 by SMg (Table S2). Of these, 147 (78·6%) were detected by both methods, 40 (21·4%) were identified only by SoC, and 28 (14·9%) were detected only by SMg.

We next examined whether performance varied by genus (Fig. 2(c)). Some taxa were consistently well detected, with F₁-scores of 1 for *Plasmodium* (n = 13), *Cryptococcus* (n = 5), *Enterocytozoon* (n = 6), and *Taenia* (n = 5). High but slightly reduced agreement was observed for *Entamoeba* (F₁ = 0·88, n = 19), *Endolimax* (F₁ = 0·91, n = 6), and *Pneumocystis* (F₁ = 0·92, n = 7). Lower performance was observed for Saccharomycotina (F₁ = 0·85, n = 50), *Aspergillus* (F₁ = 0·80, n = 15), *Blastocystis* (F₁ = 0·79, n = 17), and *Giardia* (F₁ = 0·53, n = 11). Some taxa that were rarely detected were consistently missed by SMg, including *Dicrocoelium*, *Strongyloides*, *Mansonella*, and *Trichuris*.

### Shotgun metagenomics performance varies with the sample type

Overall, SMg showed strong concordance with SoC at the genus-level, achieving a global pooled F₁-score of 0·84. At the species level, the global F₁-score decreased to 0·74 (Fig. S2). However, these values reflect pooled performance across the cohort. Since clinical interpretation also depends on sample type, we analyzed performance by sample type. The highest genus-level F₁-scores were observed in blood and biological fluids (F₁ = 0·98 and 0·97, respectively), followed by faeces (F₁ = 0·82) and respiratory fluids (F₁ = 0·79).

Per-sample F₁-scores were calculated (Fig. 2(d)) and reflect the proportion of expected organisms correctly identified in each sample, allowing comparison across different sample types. Performance varied significantly at both the genus (Kruskal–Wallis p < 0·001) and species (p = 0·003) levels. Blood and biological fluids achieved the highest accuracy, with a median F₁ score of 1 (interquartile range, IQR [1–1]). Pairwise testing revealed that the per-sample F₁-scores for blood and biological fluids were significantly higher than those for respiratory fluids (median 0·80 [0·00–1]; p = 0·001 and p = 0·031, respectively). Faeces showed higher variability and lower concordance than blood samples (median F_1_ = 1 [0·67–1]; p = 0·03), partly due to additional SMg detections and higher taxonomic diversity (Fig. 2(e)). Tissue biopsies yielded a median F₁ of 1 [0·60–1], with no significant differences compared to other sample types. At the species level, the median F₁ for blood remained significantly higher than that observed for respiratory fluids (1·00 [1·00–1.00] versus 0·67 [0–0·67]; p = 0·003), while other comparisons were not significant. Reduced and variable performance in respiratory fluids, faeces, and biopsies suggests that microbial load and diversity may be sample-type specific.

### Impact of microbial load on SMg detection performance

To assess the impact of microbial load on SMg performance, the quantification cycle (Cq) values (n = 57) from qPCR were compared between microorganisms detected by SMg (SMg+) and those missed by SMg (SMg−; Fig. 3(a)). The Cq values were lower for SMg+ microorganisms (median 26·5 [22·4–30·8], n = 46) than for SMg− microorganisms (median 37·0 [34·6–40·0], n = 11; Mann–Whitney *U* test, p < 0·001). No organism with a Cq value below 28·6 was missed by SMg. Cq values correlated inversely with RPM (Spearman’s *ρ* = −0·66, CI [-0·79 to −0·48], p < 0·001; Fig. 3(b)). The fitted relation was [Cq = 31·5 − 2.30 × log10(RPM)], indicating that a one-unit increase in Cq was associated with a 0·44-unit decrease in log10(RPM), *i.e*., a 2·7-fold reduction in RPM.

**Fig. 3.**
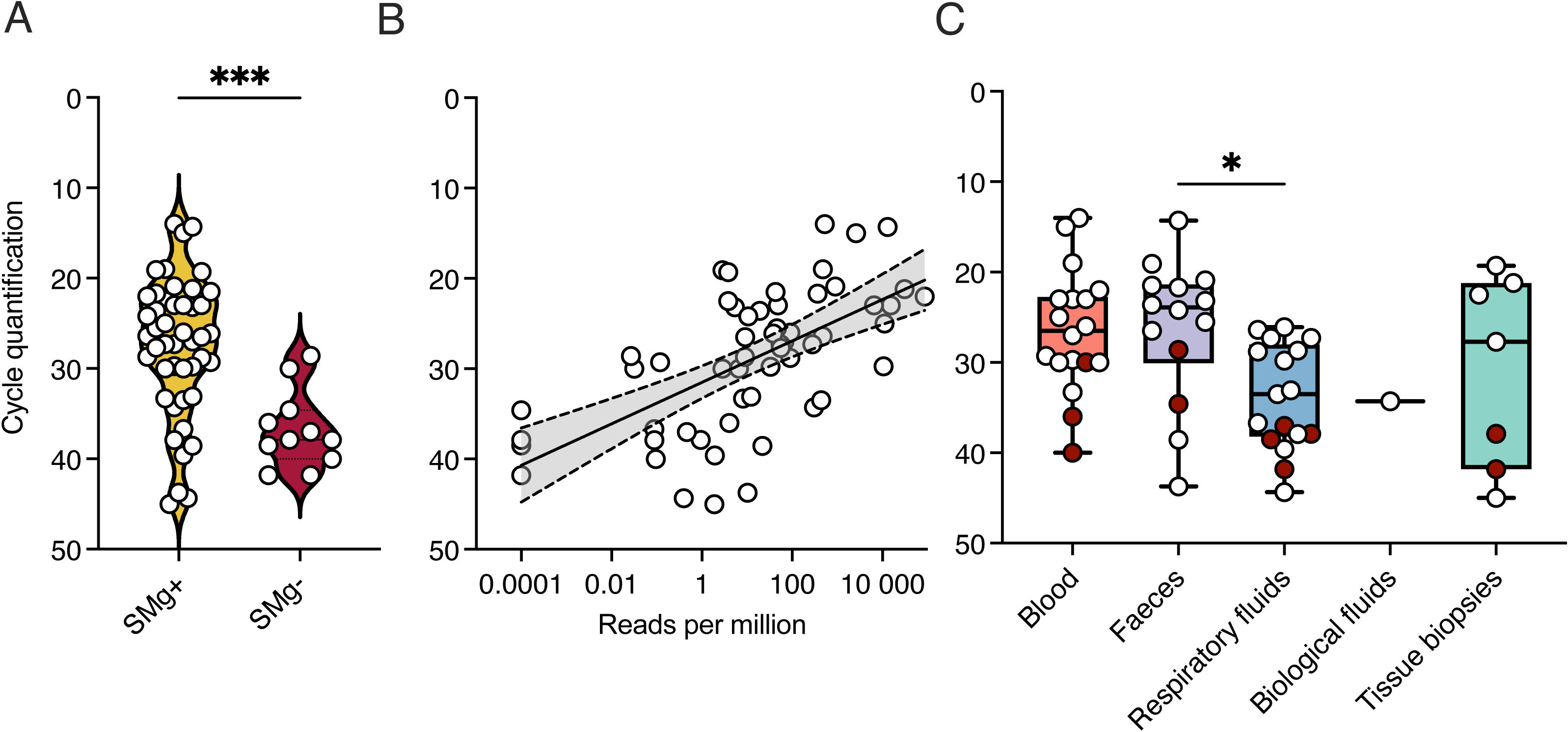
Impact of microbial load on shotgun metagenomic (SMg) performance. (a) Quantitative PCR cycle of quantification (Cq) values for microorganisms identified by SMg (SMg+) versus missed (SMg−); Mann–Whitney U test, ***p < 0.001. (b) Correlation between Cq and log₁₀-transformed reads per million (RPM) [Spearman’s ρ = −0.66 (95% CI −0.79 to −0.48), p < 0.0001]. (c) Distribution of Cq values by sample type. Red points indicate taxa not detected by SMg. Kruskal–Wallis test, p = 0.03. * Dunn’s post-hoc test, faeces vs respiratory fluids, p = 0.04.

Microbial load also differed by sample type (Kruskal–Wallis test, p = 0·03; Fig. 3(c)), with significant differences observed between faeces and respiratory fluids (Dunn’s test, p = 0·04). These results indicate that microbial load and community complexity may vary by sample type, contributing to variability in F₁-scores. To account for these sample-type effects, we next established RPM thresholds for each sample type.

### RPM thresholds for eukaryote identification by sample type

The median sequencing depth for all samples was 39,012,668 DNA reads, and 20,472,961 RNA reads (IQR [13,343,858–55,275,308] and [14,726,506–28,971,948], respectively). Thresholds that maximized Youden’s index on the ROC curve were estimated for each sample type (Fig. 4). For respiratory fluids and faeces, the optimal thresholds were 0·07 RPM (area under curve (AUC) 0·94, sensitivity 88·9%, specificity 90·7%) and 0·06 RPM (AUC 0·91, sensitivity 84·0%, specificity 91·4%), respectively. For blood and other biological fluids, the thresholds were 0·09 RPM (AUC 0·99, sensitivity 100%, specificity 92·2%) and 0·19 RPM (AUC 0·95, sensitivity 93·3%, specificity 92·8%), respectively. For tissue biopsies, a higher threshold of 0·57 RPM was obtained (AUC 0·90, sensitivity 78·9%, specificity 97·4%). When all sample types were considered together, a combined threshold of 0·06 RPM (AUC 0·93, sensitivity 87·8%, specificity 90·7%) was obtained.

**Fig. 4.**
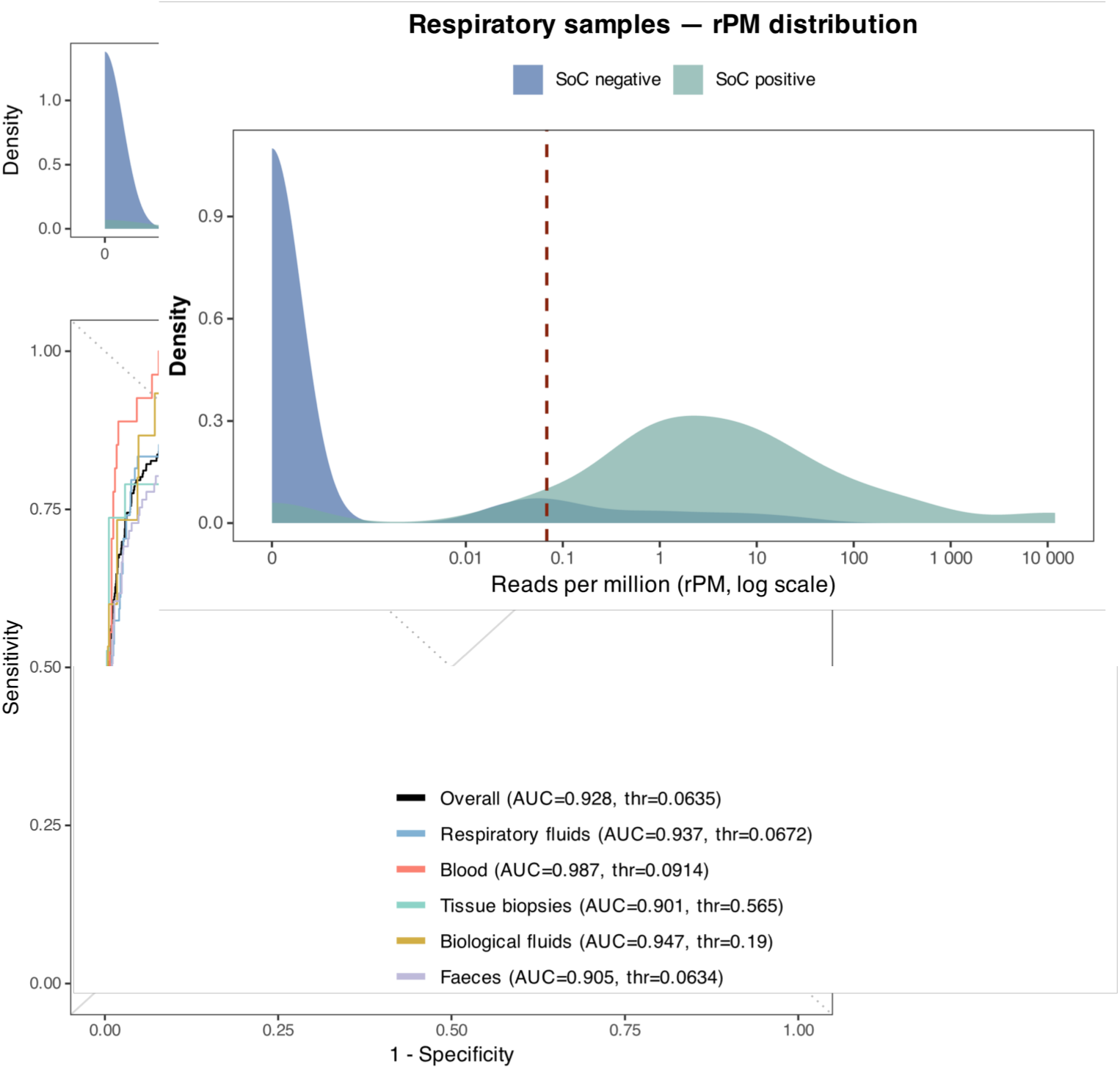
Optimal reads per million (RPM) thresholds for eukaryote detection. The upper panel displays the global density distributions of RPM for standard of care (SoC) negative (blue) and SoC positive (green) samples. The red dashed line indicates the global Youden-derived threshold (0.06 RPM). The lower panel displays receiver operating characteristic (ROC) curves stratified by sample type. The black curve represents all sample types combined (AUC 0.93, threshold 0.06 RPM). Colored curves correspond to respiratory fluids (light blue, AUC 0.94, threshold 0.07 RPM), blood (red, AUC 0.99, threshold 0.09 RPM), tissue biopsies (teal, AUC 0.90, threshold 0.57 RPM), biological fluids (gold, AUC 0.95, threshold 0.19 RPM), and faeces (purple, AUC 0.91, threshold 0.06 RPM).

The performance and RPM thresholds also varied depending on the identified taxon within the sample types (Fig. 5). *Aspergillus* was consistently detected at very low read counts in respiratory fluids (optimal threshold 0·08 RPM, AUC 0·87, sensitivity 90·0%, specificity 77·3%), whereas Saccharomycotina yeasts required a higher threshold (2·4 RPM, AUC 0·83, sensitivity 56·7%, specificity 95·5%). In tissue biopsies, the optimal threshold for the specific detection of *Aspergillus* was 1·7 RPM (AUC 1·00, 100% sensitivity and specificity). In faeces, protozoa such as *Endolimax* and *Entamoeba* were reliably detected at low RPM (optimal thresholds of 0·01 and 0·06 RPM, respectively), whereas *Blastocystis* required a higher threshold of 0·19 RPM (AUC 0·93, sensitivity 92·3%, specificity 93·5%). These findings underscore the importance of tailoring RPM thresholds to both sample types and taxa for the accurate identification of eukaryotes.

**Fig. 5.**
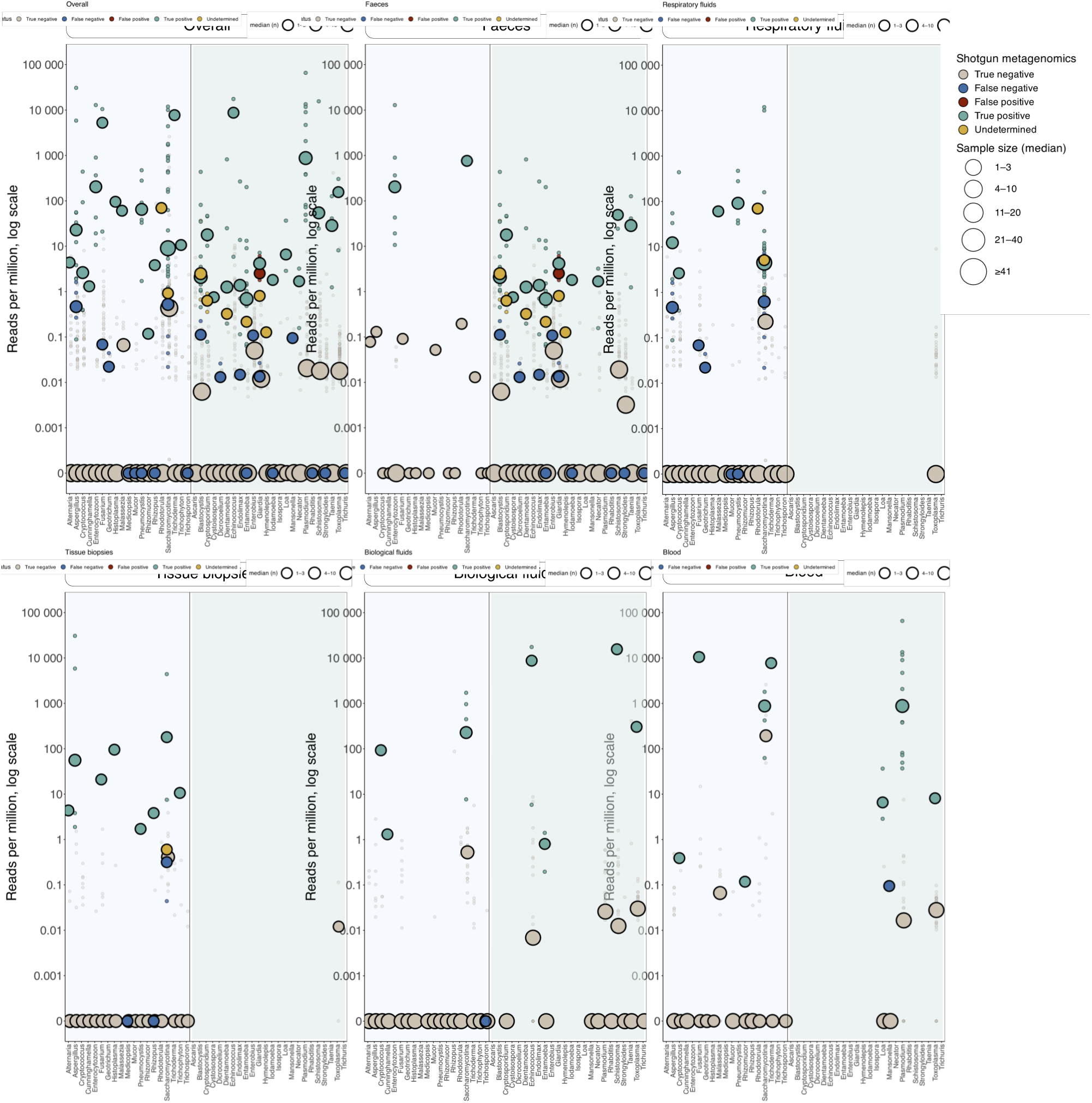
Distribution of reads per million (RPM) and concordance for eukaryotes of medical interest. Each dot corresponds to the RPM for a single microorganism–sample pair, stratified by sample type (faeces, respiratory fluids, blood, tissue biopsies, biological fluids). Concordance was assessed against the composite reference standard (standard of care plus orthogonal testing). Color codes indicate classification: true positive (green), false positive (red), false negative (blue), true negative (grey), and undetermined (gold).

## DISCUSSION

We conducted the first large-scale evaluation of clinical SMg for eukaryotic organisms. We analyzed 198 samples from five major types against the SoC diagnostics. SMg covered more than 40 genera of fungi and parasites, and showed strong genus-level agreement with SoC (F₁ = 0·84). Performance varied by sample type: faeces and respiratory fluids showed lower F₁-scores, and tissue biopsies presented the highest proportion of missed detections, likely due to sampling constraints [25]. False negatives were predominantly associated with low-abundance taxa, such as *Rhabditis*, *Strongyloides,* and *Trichuris*.

Our study has demonstrated that microbial load is a major determinant of SMg sensitivity, as indicated by its correlation with qPCR Cq values [5,26]. Higher Cq values, indicating lower pathogen abundance, were consistently associated with SMg false negatives. Therefore, several studies have proposed minimum read thresholds to ensure the reliable interpretation of low-abundance microbial signals. Miller and colleagues [27] used a composite gold standard in cerebrospinal fluid samples to validate an RPM-ratio (RPM_sample_ /RPM_NTC_) threshold of ≥10 for confident pathogen detection. Similarly, Charalampous and colleagues [28] and Alcolea-Medina and colleagues [29] suggested a minimum of five reads for the detection of fungal pathogens, such as *Aspergillus* and *Candida* species, in respiratory fluids. Other studies have applied a threshold of one pathogen-specific read for fungi, particularly *Aspergillus* [14,30]. In our study, the ROC curve analysis identified an optimal threshold of ∼0·1 RPM for eukaryotes (*i.e.*, fungi and parasites). Given a median sequencing depth of around 60 million reads per sample, this corresponded to an average of six reads per sample. While this aligns with the lower end of the thresholds reported in the literature, a cut-off point this low may be inadequate for reliably detecting pathogens, particularly in complex clinical samples or when dealing with low-abundance organisms. Although low thresholds (0·1 RPM) provide high sensitivity, they can lead to ambiguous interpretation when the number of reads detected is very low. Guaranteeing 10 pathogen-specific reads may require sequencing depths of ∼100 million reads per sample. In addition, it may be necessary to adapt sequencing strategies to exceed the 10 million reads often used for bacteria and viruses [18,23]. Currently, sequencing accounts for less than 30% of SMg costs, and this price continues to decline as high-throughput platforms are adopted. For laboratories implementing highly sensitive, pan-pathogen metagenomic diagnostics, a target of around 100 million reads per sample seems reasonable and cost-effective, as it improves detection without substantially increasing costs. Additionally, RPM thresholds need to be adjusted according to both sample type and taxon. However, for rare taxa and underrepresented sample types, larger cohorts will be required to derive robust and generalizable thresholds.

Importantly, detection does not necessarily imply pathogenicity or clinical causality, particularly in sample types with complex microbiota or in the context of colonization or residual DNA from non-viable organisms. Consequently, SMg results should be interpreted in conjunction with clinical, radiological, and biological data. In this study, comparisons were performed against standard of care assays, which are also based on detection methods, allowing for the evaluation of analytical concordance rather than a direct assessment of clinical infection.

To conclude, our evaluation of SMg’s analytical performance in detecting fungi and parasites across a wide range of clinical samples demonstrated a strong correlation with SoC testing. Using the strictest threshold (0·1 RPM) for all sample types is a practical way to maintain sensitivity when working with challenging samples. SMg is a valuable second-line tool that complements conventional methods, improving patient care for complex, hypothesis-free diagnostics. It is particularly useful in the fields of mycology and parasitology, where rare, unexpected or travel-related pathogens are often encountered, sometimes dating back many years.

## ACKNOWLEDGMENTS

We thank AP-HP (Assistance Publique–Hôpitaux de Paris) and Université Paris-Est Créteil for institutional support. We gratefully acknowledge Christel Cabon and Florent Guillemet for their technical support. The data were obtained as part of routine work at the University Hospital of Créteil, Créteil, France. The graphical abstract was created using BioRender.com.

## Declaration of Generative AI and AI-assisted technologies in the writing process

All content was written by the author(s). Spelling corrections, English grammar, and turns of phrase were revised with the help of ChatGPT-5 (OpenAI), DeepL translator, and a local version of Qwen3 running on Ollama. The authors take full responsibility for the final content of the publication.

